# The role of substrate type, moisture and temperature on the vertical growth of terricolous lichens

**DOI:** 10.1101/466227

**Authors:** Julián Monge-Nájera

## Abstract

Lichens are traditionally divided into short “crustose”, intermediate “foliose” and tall “fruticose” types, a practice that hides a growth continuum. Substrate, temperature and water are thought to affect vertical growth, but such factors are difficult to measure, because, for example, the water actually available to lichens does not match rainfall patterns or even ground water levels. To reliably assess the effect of those factors, I recorded temperature, moisture, and substrate in and under individual terricolous lichen colonies in 60 fixed quadrats on April, August, October, and December of 2015 (Cerro de la Muerte, Costa Rica, 9°33′N; 83°45′W). The measurements were taken inside the colonies themselves (rather than on the general environment), covering an annual cycle of the relatively simple páramo habitat, where animals and vegetation have less impact than in lower ecosystems. The hypotheses were that lichens would grow taller on softer, warmer, and moister ground; on the Caribbean versant; and on the rainy season. Results matched the hypotheses, with one exception: lichens on soft ground were not taller than those on rock. Caribbean colonies were, on the average, 7 cm taller than those on the drier Pacific versant. Physiologically available water seems to be the main determinant of lichen vertical growth: more water means taller lichens and greater protection from climatic change for both the lichens and their microcommunities.

---

How tall can terricolous lichens grow? What determines that growth? Research to answer these two simple questions is not abundant, perhaps because lichens have traditionally been divided into short “crustose”, intermediate “foliose” and tall “fruticose” growth forms, a practice that artificially hides a growth continuum (Grube & Hawksworth, 2007; Tehler & Irestedt, 2007).

But even though we need to go beyond growth forms to better identify growth factors, it must be borne in mind that they are useful ecological concepts when researchers want to generalize and predict: for example, crustose lichens can colonize harsher habitats (Rogers, 1990) while foliose and fruticose species are good competitors in less demanding environments (Armstrong, 1993; Armstrong & Welch, 2007). In tropical latitudes, the lower temperatures and higher photosynthetic rates of mid altitudes favor foliose and fruticose species (Ceron & Quintero, 2009), but only crustose species can grow in the more demanding conditions of high altitudes (Rai, Khare, Baniya, Upreti, & Gupta, 2015).

A possible answer to the question of what is the key factor in how tall lichens can grow is the amount of water in their habitat (Rogers, 2006), but the relationship between lichens and water is complex and hard to measure, because the water available to them does not faithfully match rainfall patterns. There are two reasons for this: one related to dew and another related to the ground itself.

Recent work found that most lichens are more directly affected by dew than by rain, because dew is a key driver of annual C-assimilation, and rainfall and dew do not necessarily match (Gauslaa, 2014). Furthermore, lichens in apparently dry areas may actually get significant moisture from the underground. Temperature and ground hardness are also drivers of vegetation growth that further complicate any conclusions in field work, so there has long been a need for studies that bypass those problems (Rogers, 2006; Gauslaa, 2014).

To reach the goal of measuring the vertical growth of lichens, while at the same time obtaining reliable measurements of moisture, temperature, and substrate type that directly affect them, I present here moisture and temperature measurements taken directly inside the lichens themselves, and in the ground under them, in a relatively simple ecosystem, the Costa Rican neotropical páramo. To prevent unwarranted generalizations from a small sample or a brief period, this study is based on hundreds of individual measurements, and covers a full yearly cycle. I chose the páramo because it is a relatively simple habitat, where animals or vegetation have less impact on lichens than in lower ecosystems (see Kappelle, & Horn, 2005, 2016).

## Methods

Along 2015 I obtained datasets in Cerro Bueno Vista (better known as Cerro de la Muerte), Costa Rica, on both versants, Pacific (9°33′33.62″N - 83°45′20.78″W) and Caribbean (9°34′40.83″N - 83°45′6.78″W), from a total of 60 PVC frame quadrats placed at random locations (as recommended for the study of small scale factors in lichens: Root & McCune, 2012). The number of measurement sets was as follows: 66 in April, 94 in August, 95 in October, and 95 in December (totals: 220 measurements on the Caribbean and measurements on the Pacific versant). It was not possible to sample in June because ….

For each measurement I identified the largest lichens inside the plot and recorded data from one to three lichens, as time permitted. Measurement sets: for each lichen I recorded growth type, height over ground (with a ruler), substrate, temperature, and moisture (electronic Dataloger HOBO U-Shuttle, ONSET, Model: U-DT-1). Measurements were done by inserting the probe in the center of the lichen and, a second later, under the lichen (touching the substrate).

Additionally, on every visit I took high definition digital photographs of all plots and the photographs are freely available for corroboration: https://doi.org/10.5281/zenodo.1296967. The raw data for this study are also freely available online for critical review and reanalysis: https://doi.org/10.5281/zenodo.1296956.

### Hypotheses

1. Lichens on hard ground grow shorter because conditions are harsher (Clair, Johansen, & Rushforth, 1993)
2. Lichens on warmer ground grow higher because of faster physiological reactions (Benedict, 1990).
3. Lichens on wetter ground grow higher (Rogers, 2006).
4. Caribbean versant lichens grow higher than Pacific versant lichens because they receive more moisture (Herrera, 1985).

## Results

The substrate was soft (i.e. dirt and organic matter) in 360 measurements and hard (i.e. rock) only in 33 measurements; there were no statistical differences in moisture, temperature, or lichen height, between soft and hard soil (Kruskal-Wallis test, p> 0.05 for all variable combinations).

Crustose lichens grow less than a millimeter and their height cannot be measured with a ruler, but the mean height of foliose lichens was 15 mm (Standard Error 1.0 mm), and for fruticose it was 82 mm (Standard Error 7.0 mm).

There were 139 measurements in which the largest lichen was fruticose, 123 where it was crustacean and 107 where it was foliose, i.e. none of the growth forms dominated numerically the random quadrats (Chi 4.16, p=0.1249).

There were 13 identifiable species in the Caribbean versant and 11 identifiable species in the Pacific versant (Table 1).

**TABLE 1.**
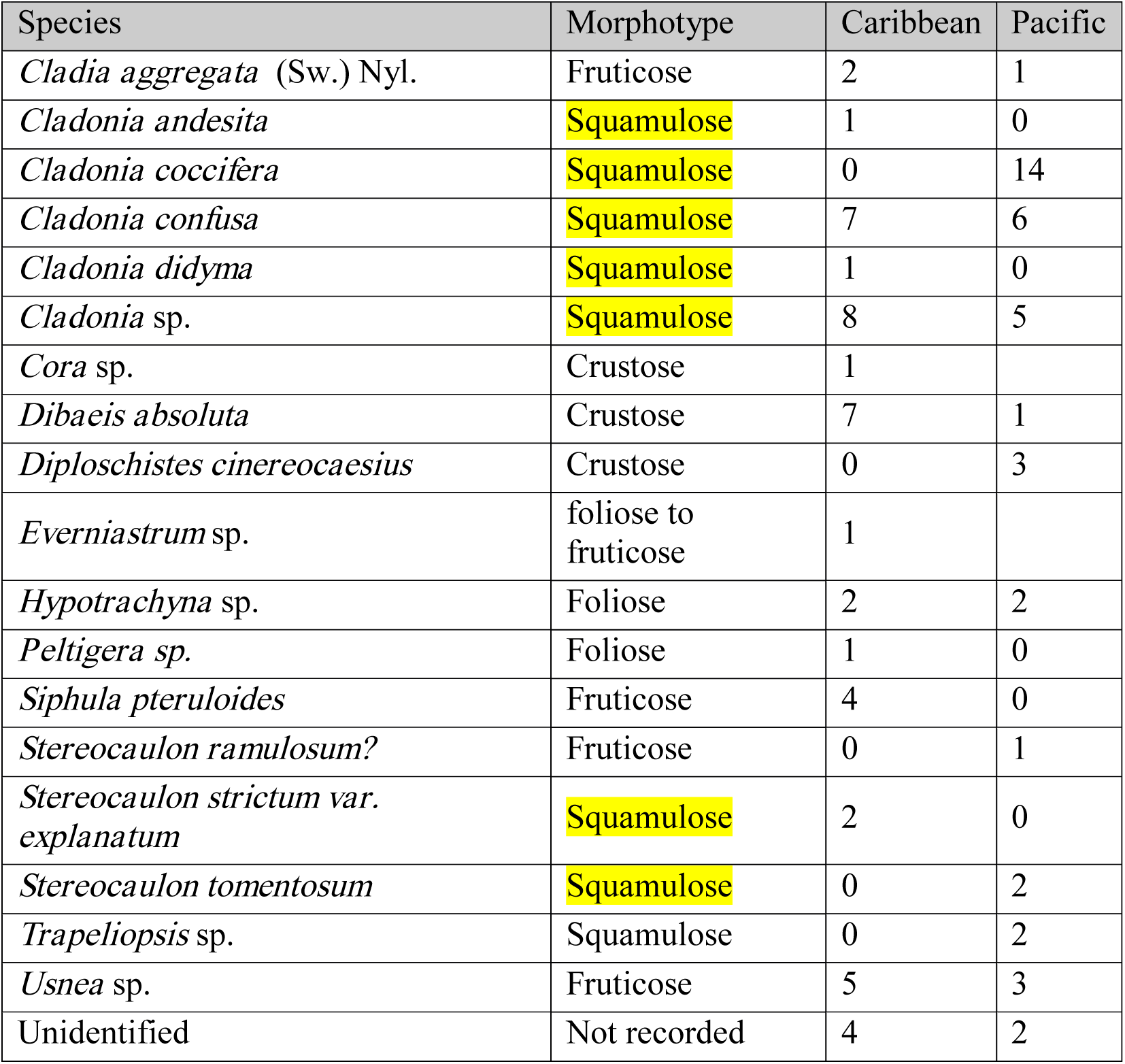
Lichen species versus versant, in number of random quadrats that had the lichen species.

In the next sections, figures are presented only when there were statistically significant patterns.

### Seasonal Pattern

in December the “Christmas Winds” reduce cloud cover, producing warmer days, colder nights, and greater temperature fluctuations in the páramo. These harsher conditions extend to early February, coinciding with higher temperatures in both soil and lichens, and matched a reduction in the mean height of the lichen colonies (Kruskal-Wallis ANOVA, p>0.05; Fig. 1).

**Fig. 1.**
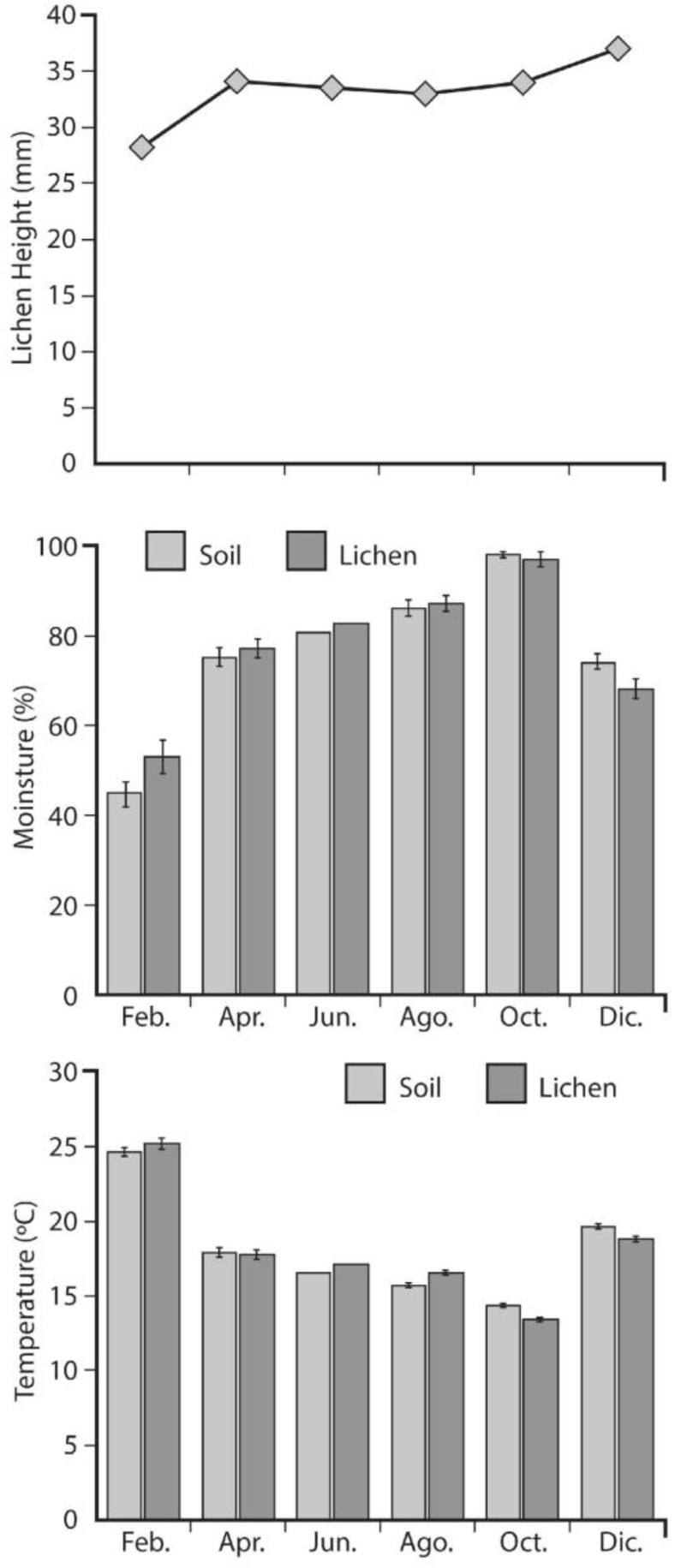
Yearly trends of mean temperature, moisture and lichen height in the Cerro de la Muerte páramo. Error bars represent the standard error of the mean (June values are interpolations because there was no sampling in that month, thus they have no error bars).

On the other hand, from April to October, when cloud cover is denser, the soil and the lichens are moister and cooler; and lichen colonies are taller than at the beginning of the year (Fig. 1).

### Effect of versant

there are no significant versant differences in temperature, but there is an important difference in moisture. In the Caribbean versant, which is wetter, both soil and lichens are moister and lichen cover is almost twice as tall as in the Pacific (Fig. 2, Kruskal-Wallis p<0.001 in all cases).

**Fig. 2.**
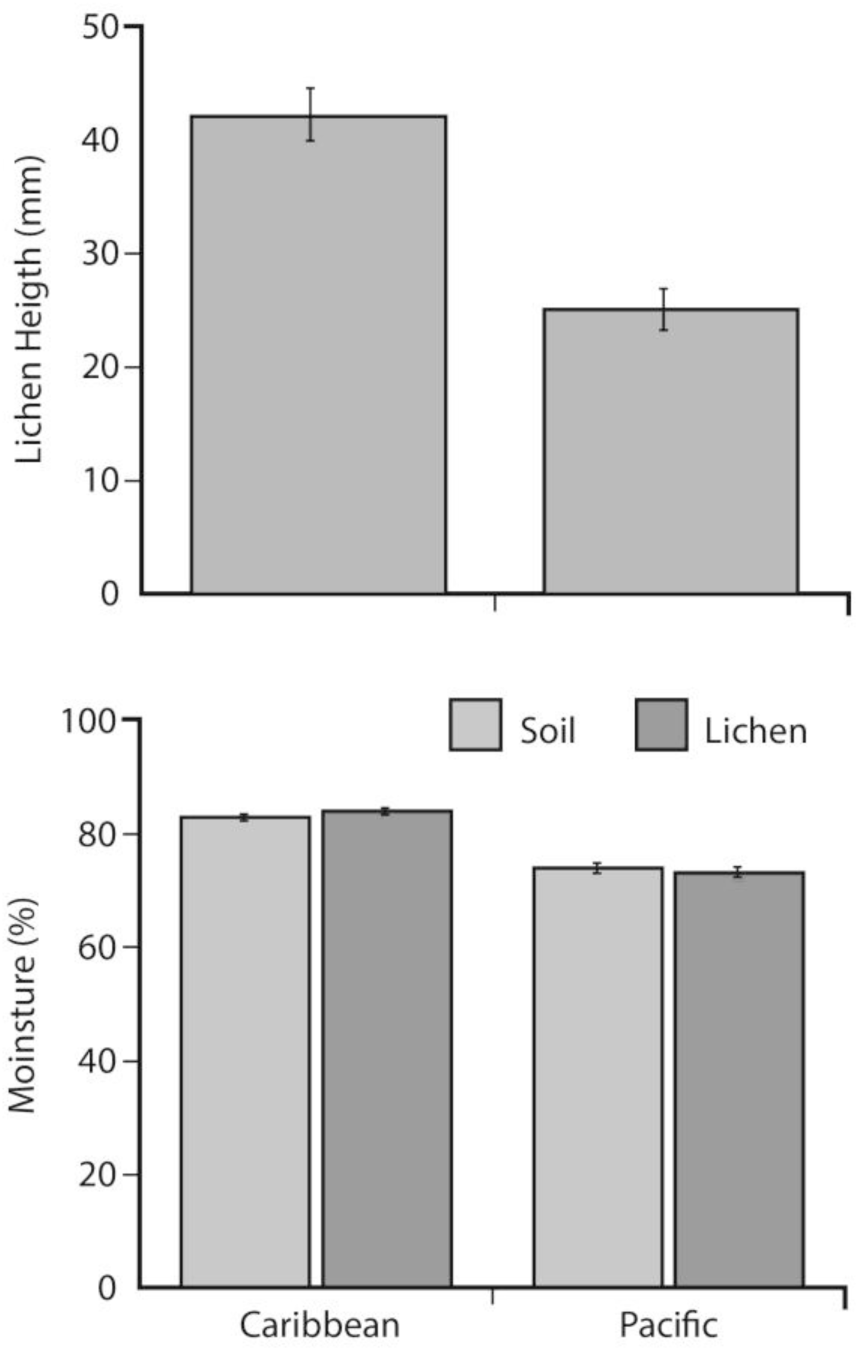
Moisture of soil and lichens, and lichen colony height, in the Cerro de la Muerte páramo (Caribbean versus Pacific versants; error bars represent the standard error of the mean).

### Growth form

in this páramo, crustose lichens provide a little cover for the soil, which matches closely the temperature and moisture of the lichen over it; in contrast, foliose and fruticose lichens keep the ground under them cooler and moister (Fig. 3), probably also providing better conditions to the associated fauna. Why do foliose lichens, which are not as tall of fruticose lichens, keep better moisture and temperature conditions? Probably because their horizontal growth better covers the ground (Cao, Zhang, Zheng, et al., 2015).

**Fig. 3.**
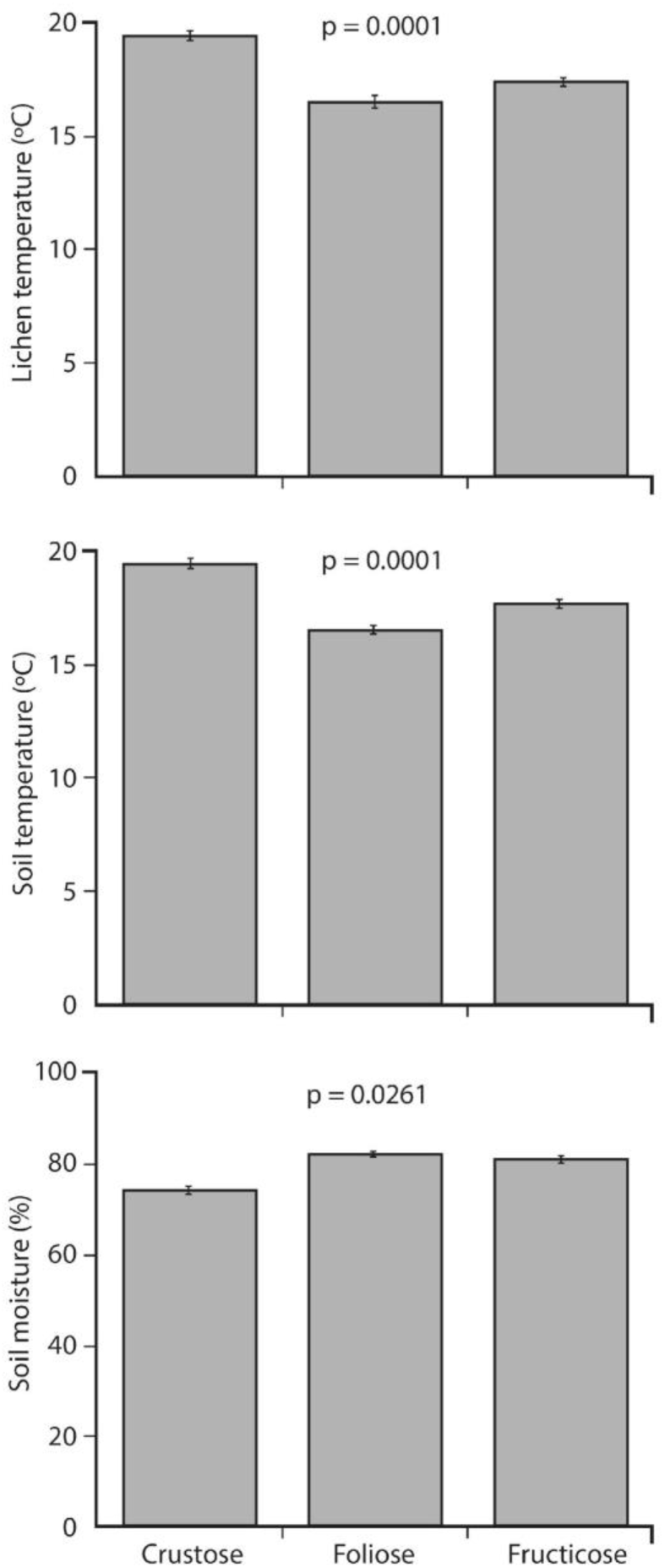
Temperature (soil and lichens) and soil moisture for each lichen growth form, in the Cerro de la Muerte páramo (Caribbean versus Pacific versants; error bars represent the standard error of the mean).

## Discussion

The lichenoflora of the Costa Rican páramos, which differs markedly from that found in lower parts of the country, has about 200 described species; these species have wide geographic ranges that cover many of the neotropical highlands, possibly as a result of geographic páramo continuity during the Pleistocene (Sipman, 2005; Kappelle & Horn, 2005, 2016). In Costa Rica, the most frequent páramo lichens are foliose and fruticose species, i.e. species that rise well above the surface (Sipman, 2005), but in this study, the number of random quadrats with each lichen type were similar. This discrepancy warrants future study.

The results are in agreement with my hypotheses, with the exception of the hypothesis that lichens grow shorter on rocky substrate. In contrast with softer soil, rock-hard substrates fluctuate more abruptly in temperature and do not absorb water as well, so I expected less vertical lichen growth in them (see Clair et al., 1993; Giordani, Incerti, Rizzi, Rellini, Nimis, & Modenesi, 2014). Perhaps the differences were not marked enough in this sample because there were proportionally few cases of hard substrate, or perhaps such generalizations do not apply to this páramo soil, but this will have to be solved by future research.

The finding that lichens grew taller in the wetter versant (Caribbean) and that the mean height of the colonies is lower after the short dry period at the end of the year, agree with previous findings from temperate regions (e.g. xxxxx). Good levels of hydration allow optimal levels of net carbon dioxide fixation (Grube & Hawksworth, 2007) and allow faster growth (Merinero, Martínez, Rubio-Salcedo, & Gauslaa, 2015). This is precisely the advantage of the Caribbean versant because Trade Winds introduce moisture from the Caribbean sea, while highlands block the arrival of clouds to the Pacific side (Herrera, 1985).

The finding that shorter lichens (i.e. crustose species) basically have the same temperature and moisture as the substrate is not surprising. There are no known equivalent tropical data, but Cao, Zhang, Zheng, Liu, & Zhou (2015) reported similar results in Antarctic lichens that showing species with a tufted thallus were better protected from external conditions than those with low lying thalli.

Terricolous lichens harbor important invertebrate communities (Bokhorst, Asplund, Kardol, & Wardle, 2015), and the more favorable microclimatic conditions of foliose and fruticose species in the páramo must be important to the many species that live in them. Besides tardigrades, little if anything is known about the microfauna of Costa Rican páramo lichens (Mehlen, 1969).

The implication of these findings is that organisms living in the crustose species will be the most affected by global warming, because those lichens offer little microclimatic protection (see Bokhorst et al., 2015). This negative effect is also expected to affect the lichens themselves (Bässler, Cadotte, Beudert, Heibl, Blaschke, Bradtka, & Müller, 2015) as well as the microbial communities under them (Maestre, Escolar, Bardgett, Dungait, Gozalo, & Ochoa, 2015).

This closer look at lichens, on the other hand, suggests that physiologically available water is the main determinant of lichen vertical growth, and that more water means taller lichens and greater protection from climatic change for both the lichens and their microcommunities.

## Acknowledgements

I thank Maribel Zúñiga, Frank González and Sergio Quesada for their valuable field work and Zaidett Barrientos for financial and logistical support, and for useful comments to improve the manuscript. Carolina Seas and Ligia Bermúdez helped greatly with data analysis and figures. I also thank the Servicio de Parques Nacionales for supporting this study. The project was financed by Vicerrectoría de Investigación UNED de Costa Rica (*Páramos neotropicales: amenazas al ecosistema por cambio global. Etapa 1*). Harrie J.M. Sipman (Botanischer Garten und Botanisches Museum, Berlin) identified the lichens and gave valuable comments to improve the manuscript.

## Resumen

Según su altura, tradicionalmente, los líquenes se dividen en “crustáceos” “foliosos” y “fruticulosos”, pero estas categorías artificiales ocultan un *continuum* de crecimiento. Se cree que la temperatura, el sustrato y el agua definen el crecimiento vertical de los líquenes del suelo, pero estos factores son difíciles de medir, porque, por ejemplo, el agua realmente disponible para los líquenes no siempre coincide con la lluvia o el agua en el suelo. Para evaluar de forma fiable el efecto de estos factores, medí temperatura, humedad y sustrato en líquenes de 60 cuadrículas aleatorias en abril, agosto, octubre y diciembre de 2015 (Cerro de la Muerte, Costa Rica, 9°33′N; 83°45′W). Las mediciones se tomaron en el interior de los líquenes (no en el ambiente externo) durante un ciclo anual en el relativamente simple ecosistema del páramo, donde los animales y la vegetación tienen menos impacto en los líquenes que en la bajura. Las hipótesis eran que los líquenes crecerían más en suelo suave, más cálido y húmedo; en el lado del Caribe; y en la época de lluvias. Los resultados coinciden las hipótesis, con una excepción: los líquenes de suelo suave no crecieron más que los de suelo rocoso. Del lado caribeño los líquenes fueron 7 cm más altos que los del Pacífico, que es más seco. El agua fisiológicamente disponible parece ser el principal determinante del crecimiento vertical de los líquenes: más agua significa líquenes más altos y una mayor protección contra el cambio climático, tanto para los líquenes como para sus microcomunidades.

### Palabras clave

variables físicas, clima, líquenes, alturas, efecto del cambio climático.

